# Molecular and transcriptional structure of the petal and leaf circadian clock in *Petunia hybrida*

**DOI:** 10.1101/641639

**Authors:** Marta I. Terry, Marta Carrera-Alesina, Julia Weiss, Marcos Egea-Cortines

## Abstract

The plant circadian clock coordinates environmental signals with internal processes. We characterized the genomic and transcriptomic structure of the *Petunia hybrida* W115 clock in leaves and petals. We found three levels of evolutionary differences. First, *PSEUDO-RESPONSE REGULATORS PhPRR5a, PhPRR5b, PhPRR7a, PhPRR7b*, and *GIGANTEA PhGI1* and *PhGI2*, differed in gene structure including exon number and deletions including the CCT domain of the PRR family. Second, leaves showed preferential day expression while petals tended to display night expression. Under continuous dark, most genes were delayed in leaves and petals. Importantly, photoperiod sensitivity of gene expression was tissue specific as *TIMING OF CAB EXPRESSION PhNTOC1* was affected in leaves but not in petals, and *PhPRR5b, PhPRR7b* and the *ZEITLUPE* ortholog *CHANEL, PhCHL*, were modified in petals but not leaves. Third, we identified a strong transcriptional noise at different times of the day, and high robustness at dawn in leaves and dusk in petals, coinciding with the coordination of photosynthesis and scent emission. Our results indicate multilayered evolution of the *Petunia* clock including gene structure, number of genes and transcription patterns. The major transcriptional reprogramming of the clock in petals, with night expression may be involved in controlling scent emission in the dark.

**Highlight:** The petunia leaf circadian clock shows maxima during the day while petal clock does it during the night. Reaction to dark is organ specific.

## Introduction

Organisms, from bacteria to human beings, are subjected to periodic oscillations in the environment due the planet rotation around its axis. Circadian clocks are a complex set of genes allowing organisms to anticipate and adapt to daily environmental variations. In plants, the circadian clock is a network of interlocked loops comprising transcriptional, translational and posttranslational coordination (Harmer, 2009). Circadian processes have been studied in plants for a long period of time (see McClung for a historical overview, (McClung CR, 2006)). Most molecular studies have been done in *Arabidopsis thaliana*. The Arabidopsis core clock is formed by several genes. Two MYB transcription factors *CIRCADIAN CLOCK ASSOCIATED 1 (CCA1), LATE ELONGATED HYPOCOTYL (LHY)* and the *PSEUDO RESPONSE REGULATOR TIMING OF CAB EXPRESSION (TOC1*) form the so-called core clock. Later studies found other clock components including the *PSEUDO-RESPONSE REGULATOR* gene family (*PRR*), out of which PRR3, PRR5, PRR7 and PRR9 are clock genes, and the Evening Complex (EC), which is formed by the *EARLY FLOWERING 3 (ELF3), EARLY FLOWERING 4 (ELF4)* and *LUX ARRHYTMO (LUX)* proteins. In addition, other genes playing a key role and considered part of the clock include the protein with blue light reception capacity ZEITLUPE *(ZTL)* and the single copy gene *GIGANTEA (GI)*. The various models developed are based on mutually repressing genes and a set of activating genes coded by the *REVEILLE* MYB transcription factors (Hsu *et al.*, 2013). Every new discover has added a level of complexity and new interpretation of the circadian clock model (Hernando *et al.*, 2017).

Two aspects emerge from comparative genomics with lower organisms and within higher plants. First the core clock components identified in the picoeukaryote *Ostreococcus* comprise a *MYB* gene homolog to *LHY* and a *PRR* gene similar to *TOC1* (Corellou *et al.*, 2009). There is an additional blue-light receptor component with histidine kinase activity and circadian clock effects (Djouani-Tahri *et al.*, 2011). So, basic clocks maybe found with two or maybe three components that function via transcriptional control. A second aspect is that the fine tuning of the different clock modules is based to a large extent on protein-protein interactions. As protein complexes require certain stoichiometries to maintain their function they are target of genetic constraints in terms of gene dosages and are especially sensitive to gene duplications. Duplicated genes follow four paths including gene loss, maintenance of redundancy, subfunctionalization or neofunctionalization (Airoldi and Davies, 2012). Plant genomes have been subject to genome duplications and, in some cases, followed by non-random elimination of duplicated genes (Adams and Wendel, 2005; Wendel *et al.*, 2016). In *Brassica*, polyploidization events have involved subsequent gene loss but with a preferential retention of circadian clock genes as compared to house-keeping genes, supporting a gene dosage sensitivity model (Lou *et al.*, 2012).

The genomes of the garden petunia and its ancestors *Petunia axillaris* and *P. integrifolia* have been recently sequenced (Bombarely *et al.*, 2016). Petunia forms an early branching in the Solanaceae clade departing from *Solanum lycopersicon, S. tuberosum, Nicotiana spp.* and *Capsicum spp.* that have a chromosome number of n=12. Petunia has n=7 and this, together with a high activity of transposition, may have shaped a somewhat different genome evolution. Petunia shares a paleohexaplodization specific to the Solanaceae. A comprehensive analysis of the circadian clock genes found in the *Petunia* genomes shows that there is a set of genes that has remained as single copy. These include the petunia orthologs for *PRR9, PRR3, TOC1* and *LHY*. In contrast, other genes are present in two to four copies, *PRR7, PRR5, GI, ELF3* or *ELF4* (Bombarely *et al.*, 2016). Altogether these data indicate a possible departure of the circadian clock network from the one known in Arabidopsis, and suggests the evolution of the clock at different levels including gene structure, expression pattern and genetic functions.

The bulk of work on plant circadian rhythms has been done in Arabidopsis using leaf tissue and seedlings. Like in animals, there is important evidence that the circadian clock expression network differs between different organs. The current view is that the shoot apical meristem may work as a center of coordination (Takahashi *et al.*, 2015), and leaves and roots differ in the regulatory network, as a result of differences in light inputs (James *et al.*, 2008; Bordage *et al.*, 2016).

Petal development starts with the activation of the so-called B function genes in both gymnosperms and angiosperms (Theissen and Becker, 2004). The initial transcriptional activation is followed at early stages by an autoregulatory positive regulation of the MADS-box genes controlling petal morphogenesis in *Antirrhinum*, Arabidopsis and petunia (Schwarz-Sommer *et al.*, 1992; Goto and Meyerowitz, 1994; Jack *et al.*, 1994; Zachgo *et al.*, 1995; Samach *et al.*, 1997; Vandenbussche *et al.*, 2004). Once organ identity is established and right after anthesis, there is a transcriptional reprogramming (Manchado-Rojo *et al.*, 2012). Furthermore, in sympetalous flowers with petals forming a tube and a limb, both parts of the flower appear to have different functions and transcriptional control (Delgado-Benarroch *et al.*, 2009; Manchado-Rojo *et al.*, 2014). The petal function after anthesis includes concealing the sexual organs and attracting pollinators. The lifespan of a flower is relatively short with most flowers surviving two to five days after anthesis. After anthesis, metabolism and scent emission changes rapidly (Muhlemann *et al.*, 2012; Weiss *et al.*, 2016). Flowers enter rapid senescence upon pollination as a result of ethylene release (Shaw *et al.*, 2002; van Doorn and Woltering, 2008; Liu *et al.*, 2011).

Floral scent release depends on petal development in a quantitative way (Manchado-Rojo *et al.*, 2012), and is circadian regulated in monocots and dicots such as *Antirrhinum, Narcissus*, rose or petunia (Helsper *et al.*, 1998; Kolosova *et al.*, 2001; Verdonk *et al.*, 2003; Hoballah *et al.*, 2005; Ruíz-Ramón *et al.*, 2014). Most flowers analyzed emit scent preferentially during the day or during the night. The *LHY* and *ZTL* orthologs control scent emission in *Petunia* and *Nicotiana attenuata* (Fenske *et al.*, 2015; Yon *et al.*, 2015; Terry *et al.*, 2019). Both emit higher quantities during the night, indicating an identity and circadian component controlling this trait.

In the current work, we have addressed the structure of the petunia circadian clock from three different perspectives. The gene structure diverges as *PRR* paralogs have different intron numbers and *PhGI1* and *PhGI2* vary in the coding region. The transcriptional structure showed maximum expression during the day in leaves and during the dark in petals. This maximum tended to delay in both tissues under constant darkness conditions. We further identified opposite levels of transcriptional noise at dawn in leaves and dusk in petals. Our results reflect the evolution of the plant circadian clock at different overlapping levels and indicate an organ specific transcriptional structure of the plant circadian clock.

## Materials and Methods

### Plant materials and experiment design

We used the *Petunia hybrida* W115 Mitchell for all the analysis. Plants were grown in the greenhouse under natural conditions. Experiments under controlled conditions in growth chambers were performed as described (Mallona *et al.*, 2011*a*), with the following modifications. For the control experiment, plants were adapted to light:dark growth chamber conditions for at least 1 week. Day:night (12LD) conditions were matched with thermoperiods of 23 °C:18 °C during the light and dark periods. Zeitgeber time (ZT) was defined as ZT0 for light on and ZT12 for light off. In the second experiment, plants were transferred from 12LD cycle to a continuous dark cycle (12DD) with the same temperature regimes.

Flowers were marked before opening, and samples were taken at day 2-3 after anthesis. We used the petal limbs for all experimental procedures. We used young leaves with a length of 1.5-2.5 cm for all the experiments. Sampling of petal limbs and leaves was made every three hours, starting at ZT0 and tissues were immediately frozen in liquid nitrogen. In the case of 12DD experiment, sampling also started at ZT0, during the first 24h under continuous dark.

### Phylogeny and bioinformatics

Gene models of Solanaceae were obtained from (https://solgenomics.net/), *Antirrhinum* from (http://bioinfo.sibs.ac.cn/Am/) (Li *et al.*, 2019*b*), TAIR (https://www.arabidopsis.org/), Phytozome (https://phytozome.jgi.doe.gov/pz/portal.html) and NCBI (https://www.ncbi.nlm.nih.gov/). We used the corresponding predicted proteins to identify the intron-exon boundaries using Genewise (Birney *et al.*, 2004). The corresponding exon-intron boundaries were plotted using the exon-intron graphic maker (http://wormweb.org/exonintron). Protein alignment was performed with CLUSTALX (Larkin *et al.*, 2007). Phylogenetic analysis was performed with the R libraries “ape” and “phangorn” (Paradis *et al.*, 2004; Schliep, 2011) (R version 3.5.1), using the Maximum Likelihood as statistical method, JTT (Jones, Taylor and Thornton, (Jones *et al.*, 1992)) as model of amino acid substitution and 500 bootstrap replicates. Trees were visualized and annotated with “ggtree” (Yu *et al.*, 2017) using R. Protein domains were predicted using the web-based tool PROSITE (Hulo *et al.*, 2006), schematic proteins were plotted with the R package “drawProteins” (Brennan, 2018). The protein sequences used in the phylogenetic reconstruction are listed in the Supplementary Table S1 and Supplementary Table S2.

Detection of rhythmic gene expression was performed using the non-parametric statistical algorithm JTK_CYCLE (Hughes *et al.*, 2010) implemented in the R package “MetaCycle” (Wu *et al.*, 2016). We analyzed leaves and petals, under two light conditions, 12h light/12h dark (12LD) and constant darkness (12DD). Differences between two time series, were tested using an harmonic ANOVA (HANOVA) implemented in the R package “DODR” (Thaben and Westermark, 2016). We plotted the graphics with “ggplot2” (Wickman, 2017).

### Gene expression analysis by qPCR

RNA was extracted from three biological replicates per time point of leaves and corollas using acid phenol (Box *et al.*, 2011). Concentrations were measured using NanoDrop (Thermo-Fisher). Equal amounts of total RNA were used to obtain cDNA using Maxima kits (Thermo-Fisher).

PCR analysis was performed as described before (Mallona *et al.*, 2010), the following protocol was used for 40 cycles: 95 °C for 5 s, 60 °C for 20 s and 72 °C for 15 s (Clontech SYBR Green Master Mix and Mx3000P qPCR Systems, Agilent Technologies). Primers for circadian clock genes were designed using pcrEfficiency (Mallona *et al.*, 2011*b*) (Supplementary Table S3) and the following protocol was used for 40 cycles: 95 °C for 5 s, 60 °C for 20 s (55 °C for *PhGI1* and *PhGI2*) and 72 °C for 15 s. Samples were run in duplicate. Primer combinations were tested with genomic DNA from Mitchell and we found that all of them gave a single copy DNA on agarose gels. The endpoint PCR was further verified by melting point analysis where all primer combinations gave a single peak of melting (Supplementary Fig. S1). Normalized expression was calculated as described (Schmittgen and Livak, 2008) and *PhACT* was the internal control gene, a stable gene in circadian studies in petunia leaves and petals (Terry *et al.*, 2019).

## Results

### The duplicated *PRR5, PRR7* and *GI* diverge in intron number and coding sequence

We used the laboratory line *Petunia hybrida* W115, also known as Mitchell, which contains the circadian clock genes corresponding to *P. axillaris* (Bombarely *et al.*, 2016) for a detailed analysis of the structure of the *PRR* and *GI* paralogs. Several genes forming the morning and evening loops of the circadian clock in petunia have undergone gene duplication. The genome of petunia has seven *PRR* genes as *PRR7* and *PRR5* are duplicated both in *P. axillaris* and *P. integrifolia* while Arabidopsis has the canonical set of five genes, *PRR1* or *TOC1, PRR3, PRR5, PRR7* and *PRR9* involved in circadian regulation (Bombarely *et al.*, 2016). We reconstructed a phylogenetic tree of *PRR* genes of Solanaceae and Arabidopsis (Supplementary Table S1) in order to deduce the evolutionary relationships of the duplicated genes. As found previously for other Angiosperms, the *PRR* genes of Solanaceae form three major clades: the *TOC1/PRR1* clade, the *PRR7/3* clade and the *PRR9*/*5* clade (Fig. 1) (Takata *et al.*, 2010). The *PRR5a* genes of *P. axillaris, P. integrifolia* are closer to the Arabidopsis *AtPRR5* while the rest of the *PRR* genes of Solanaceae, including the *PRR5b*, form an additional subclade. This topology indicates that the *PRRa* paralogs may be an ancestral form and the *PRRb* may have been formed later and retained, in some cases as single copy genes. The *PRR7* genes also showed a similar topology where *PaxiNPRR7a* and *PinfS6PRR7a* are closer to the Arabidopsis gene than the single copy genes of the rest of the Solanaceae, and the *PRR7b* paralogs. This topology is also seen in petunia *PRR9, PRR3* and *TOC1* that are somewhat between the Arabidopsis gene and the rest of the Solanaceae, according to the early departure of *Petunia* from the rest of the family (Bombarely *et al.*, 2016).

**Fig 1.**
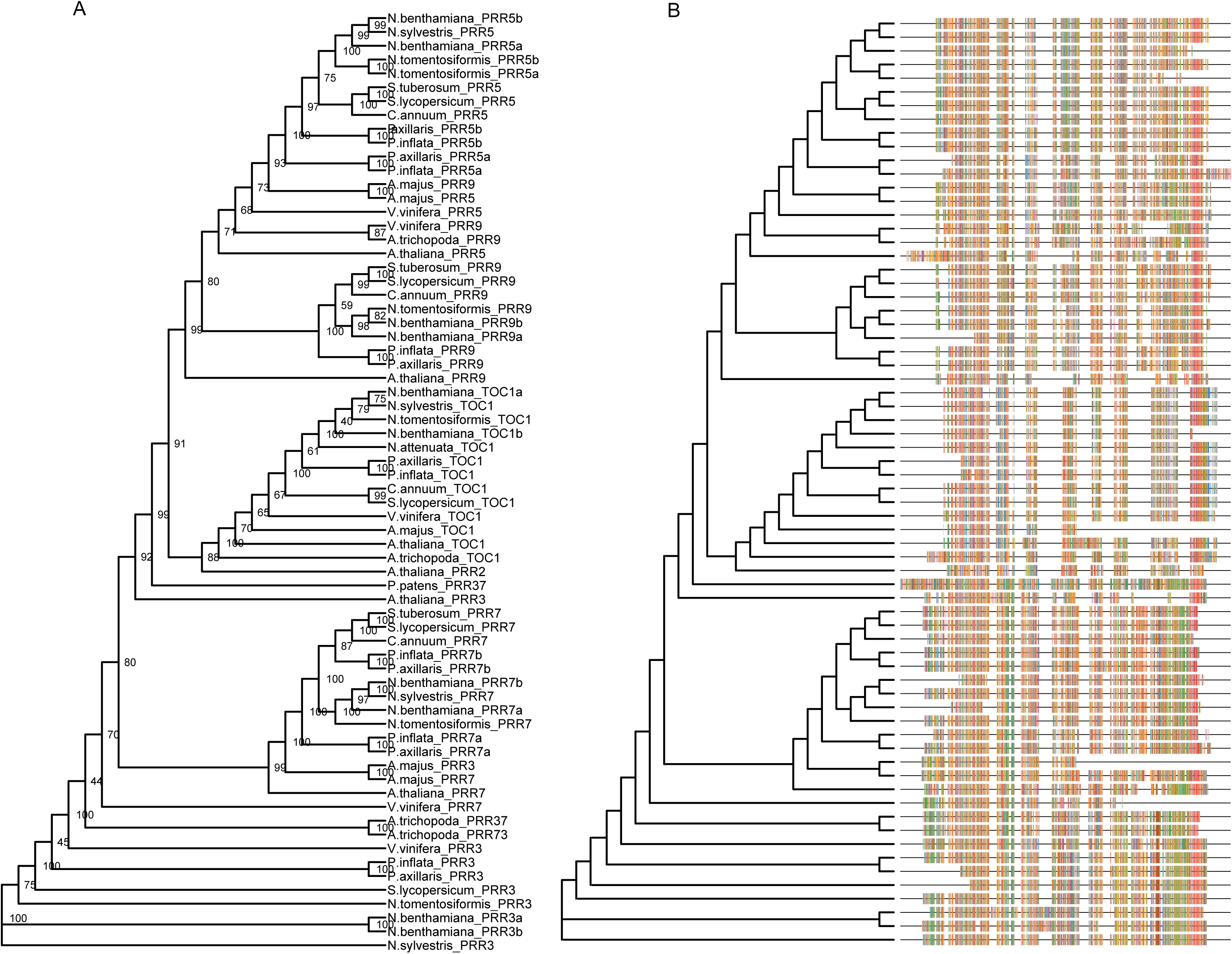
*PSEUDO-RESPONSE REGULATORS* (*PRRs*) phylogenetic tree. Amino acid sequences were aligned using CLUSTALX. Phylogenetic analysis was performed using the “ape” and “phangorn” R packages and trees were plotted with the library “ggtree” (R version 3.5.1). Initial tree was estimated using the neighbor-joining algorithm (NJ), and then phylogenetic trees were built with the Maximum Likelihood method (ML) and JTT (Jones, Taylor and Thornton) as model of amino acid substitution. The tree shows the bootstrap percentage (from 500 replicates) next to branches. The multiple sequence alignment is showed on the right-side. This tree contains 69 sequences from 14 species. Species abbreviations: A.majus (*Antirrhinum majus*), A.thaliana (*Arabidopsis thaliana*), A.trichopoda (*Amborella trichopoda*), C.annuum (*Capsicum annuum*), N.attenuata (*Nicotiana attenuata*), N.benthamiana (*Nicotiana benthamiana*), N.sylvestris (*Nicotiana sylvestris*), N.tomentosiformis (*Nicotiana tomentosiformis*), P.axillaris (*Petunia axillaris*), P.inflata (*Petunia inflata*), P.patens (*Physcomitrella patens*), S.lycopersicum (*Solanum lycopersicum*), S.tuberosum (*Solanum tuberosum*) and V.vinifera (*Vitis vinifera*). Accessions are listed in Supplementary Table S1.

We found that the gene models for *PhPRR5a* and *PhPRR5b* differ in the number of exons comprising the coding region as *PhPRR5a* has seven and *PhPRR5b* eight exons (Supplementary Fig. S2). The gene model in Arabidopsis comprises 6 exons in AT5G24470 (*AtPRR5*), indicating that changes in intron-exon structure has occurred in the evolution of the *PRR* family. The number of exons also differed between *PhPRR7a* with eight exons while *PhPRR7b* had seven exons. The Arabidopsis AT5G02810 *AtPRR7* has nine exons out of which eight correspond to coding region, thus coinciding with the phylogenetically closer *PhPRR7a*.

The PRR family of Arabidopsis has two conserved domains: REG (Response Regulatory Domain) and a CCT (CONSTANS, CONSTANS-like, and TIMING OF CAB EXPRESSION 1 [TOC1/PRR1]) (Liu *et al.*, 2016) (Supplementary Fig. S3A). We used Arabidopsis as model and we compared it with petunia sequences. We found that all the PRR members of *P. axillaris* and *P. inflata* shared the REG domain (Supplementary Fig. S3A). The CCT domain was found in all the coding genes except for PaxiNPRR7b, PinfPRR7a and PinfPRR7b. The presence of the CCT domain in PaxiNPRR7a and absence from the rest of the gene group in petunia was surprising, thus we analyzed other Solanaceae, member of the Convolvulaceae (*Cuscuta australis* and *Ipomea nil*) and Plantaginaceae (*Antirrhinum majus*). We found that the CCT domain was absent in the Solanaceae analyzed (*Capsicum annuum, C.baccatum, Nicotiana benthamiana, N.sylvestris, N.tabacum, N.tomentosiformis, Petunia axillaris, P. inflata, Solanum lycopersicum, S.melongena, S.pennellii, S.pimpinellifolium, S.tuberosum*) (Supplementary Fig. S3B). However, the CCT domain could be found in the rest of the species analyzed. This indicates an early change in the PRR7 family in Solanaceae with possible implications in clock functioning.

*GIGANTEA* is a single copy gene in the Arabidopsis genome (Fowler *et al.*, 1999) and it is found in one to three copies in the Solanaceae genomes (Bombarely *et al.*, 2016). The genes *PaxiNGI1* and *PaxiNGI2* are present in the genome of *P. hybrida* Mitchell. PhGI1, PinfS6GI1 and PinfS6GI1 share an N-terminus conserved with AtGI that was absent in PhGI2 (Fig. 2, Supplementary Fig. S4, Supplementary Table S2). Furthermore, PhGI2 has a 41 amino acid insertion that was not conserved in PinfS6GI2 or other GI genes. The PinfS6GI3 is much shorter that the other paralogs, a feature conserved in *N. benthamiana* GI3 (Fig. 2). The PinfSGI1 had an additional C-terminal fragment of 105 aminoacids absent from the rest of the GI genes analyzed (Fig. 2, Supplementary Fig. S4).

**Fig 2.**
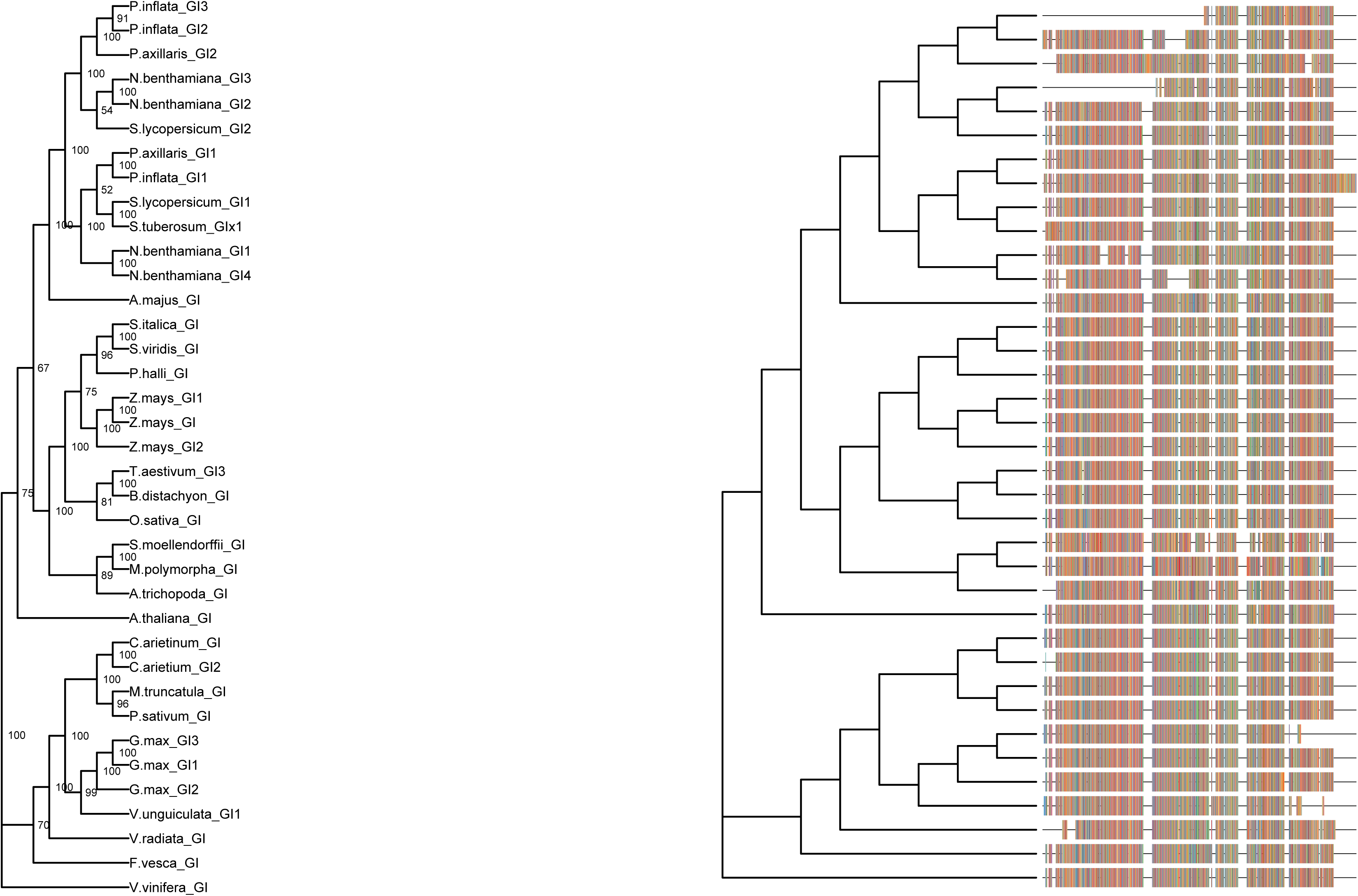
*GIGANTEA* (*GIs*) phylogenetic tree. Amino acid sequences were aligned using CLUSTALX. Phylogenetic analysis was performed using the “ape” and “phangorn” R packages and trees were plotted with the library “ggtree” (R version 3.5.1). Initial tree was estimated using the neighbor-joining algorithm (NJ), and then phylogenetic trees were built with the Maximum Likelihood method (ML) and JTT (Jones, Taylor and Thornton) as model of amino acid substitution. The tree displays the bootstrap percentage (from 500 replicates) next to branches. The multiple sequence alignment is displayed on the right-side. This tree contains 37 sequences from 25 species. Species abbreviations: A.majus (*Antirrhinum majus*), A.thaliana (*Arabidopsis thaliana*), A.trichopoda (*Amborella trichopoda*), B.distachyon (*Brachypodium distachyon*), C.arietinum (*Cicer arietinum*), F.vesca (*Fragaria vesca*), G.max (*Glycine max*), M.polymorpha (*Marchantia polymorpha*), M.truncatula (*Medicago truncatula*), N.benthamiana (*Nicotiana benthamiana*), O.sativa (*Oryza sativa*), P.axillaris (*Petunia axillaris*), P.hallii (*Panicum hallii*), P.inflata (*Petunia inflata*), P.sativum (*Pisum sativum*), S.italica (*Setaria italica*), S.lycopersicum (*Solanum lycopersicum*), S.moellendorffii (*Selaginella moellendorffii*), S.tuberosum (*Solanum tuberosum*), S.viridis (*Setaria viridis*), T.aestivum (*Triticum aestivum), S.italica (Setaria italica)*, V.radiata *(Vigna radiata)*, V.unguiculata (*Vigna unguiculata*), V.vinifera (*Vitis vinifera*) and Z.mays (*Zea mays*). Accession are listed in Supplementary Table S2.

We can conclude that the structural evolution of core circadian clock genes has occurred at several levels including changes in the number of retained paralogs, gene structure and coding region.

### The leaf clock has its maximum during the day while the petal clock shifts towards the night

The current model of the plant circadian clock defines three loops called morning, central and evening loop. These describe the time of the day when certain genes are preferentially expressed (Pokhilko *et al.*, 2012). We established the expression patterns of the different clock genes in leaves and petals. As the genes contained in *P.hybrida cv Mitchell* correspond to *P.axillaris*, we further describe them as *Ph* genes. These included the morning loop genes *PhPRR9, PhPRR7a, PhPRR7b, PhPRR5a, PhPRR5b* and *PhPRR3*. The core loop was represented by *PhTOC1* and *PhLHY*. Finally, the evening genes analyzed included *PhGI1, PhGI2, PhELF4, PhCHL* and *PhFKF*. This analysis was performed in petunia that was acclimated to light:dark conditions of 12 hour light and 12 dark (12LD) or continuous dark (12DD) conditions.

We compared three parameters between leaves and petals at 12 hours light/12 hours dark: rhythmicity of expression (oscillation), time point with maximum expression (phase) as well as amplitude, defined as is the difference between the peak or trough (maximum or minimum) and the mean value of a wave (Supplementary Table S4). Concerning the rhythmicity, most genes showed a rhythmic oscillation pattern except *PhELF4 and PhCHL* in leaves, and *PhCHL* and *PhPRR7* in petals (Supplementary Table S4).

Concerning the time of peak expression, most genes had their maximum expression during the light phase in leaves, except *PhELF4* and *PhLHY* at ZT15 and ZT21 respectively. The light phased genes peaked either during the morning at ZT4.5 (*PhPRR5a* and *PhCHL*), during midday at ZT 7.5 (*PhPRR5b, PhPRR7a, PhPRR9* and *PhTOC1)*, towards the afternoon at ZT9 *(PhGI1, PhGI2, PhPRR3* and *PhPRR7b*) or at dask at ZT 10.5 (*PhFKF*). In contrast, most of these genes shifted their expression maximum to the dark period in petals (Fig. 3) with the exception of *PhCHL, PhPRR9* and *PhPRR7a.* Among those genes that maintained their expression peak during the day or night, *PhPRR9, PhCHL* and *PhELF4* showed a delay and *PhPRR7a* an advance of 1.5 hours compared to leaves. The genes that reached their maximum during the dark period in petals could be divided in those with a peak expression early at night at ZT12 (*PhGI2* and *PhPRR7b)*, a peak towards the middle of the night at ZT 13.5 and ZT15 (*PhGI1, PhPRR3, PhPRR5a* and *PhPRR5b, PhFKF*) and those with a maximum expression at the end of the night at ZT21 (*PaxiELF4* and *PhTOC1)* (Table 1). The only gene showing a maintained expression maximum in leaves and petals was *PhLHY.*

**Table 1.**
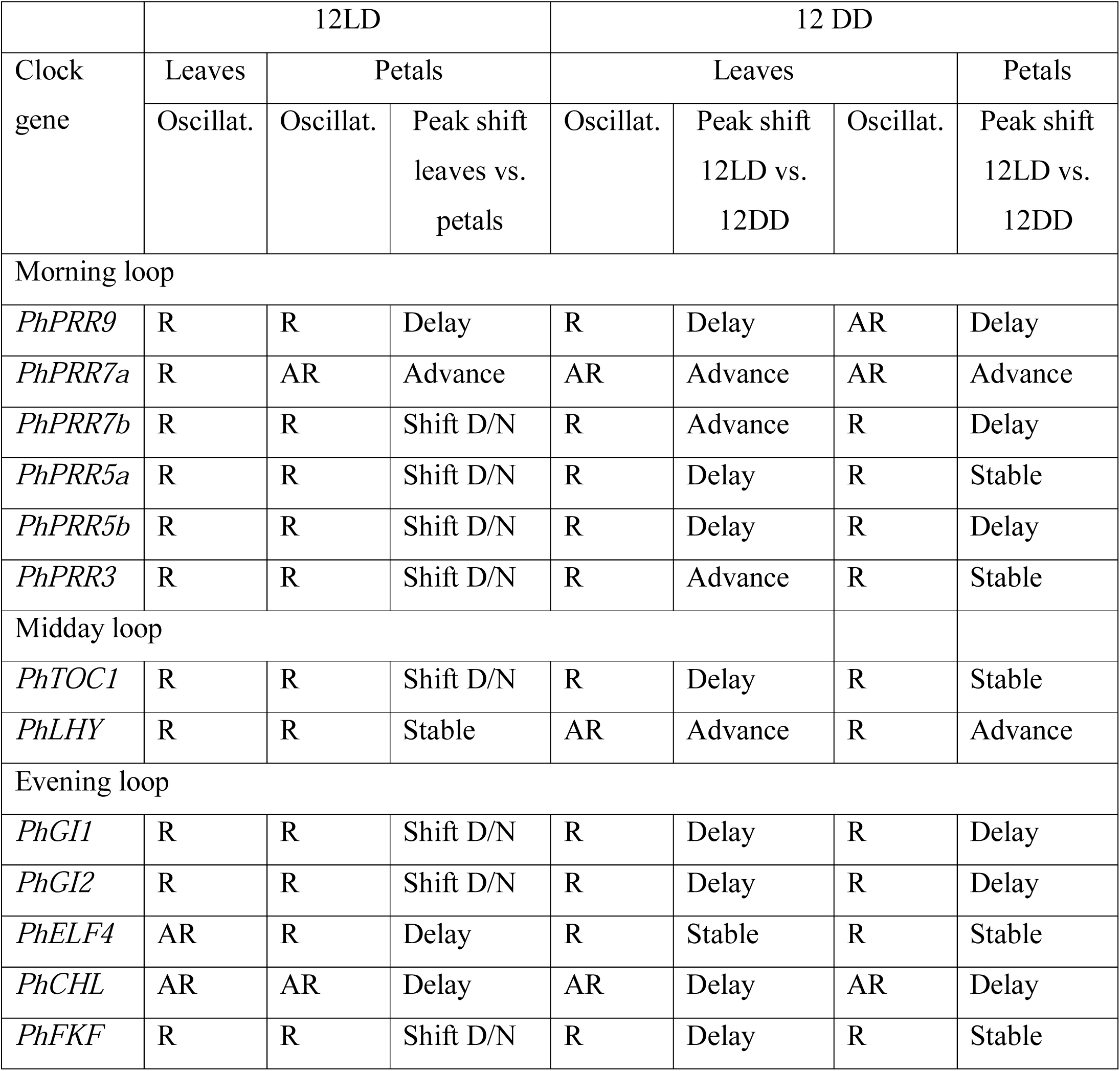
Comparison of oscillation pattern (Oscillat.) and peak shifting of morning, midday and evening loop clock genes between leaves and petals and between 12LD and 12 DD. R: Rhythmic, AR: Arrhythmic, D/N: Day-Night

**Fig 3.**
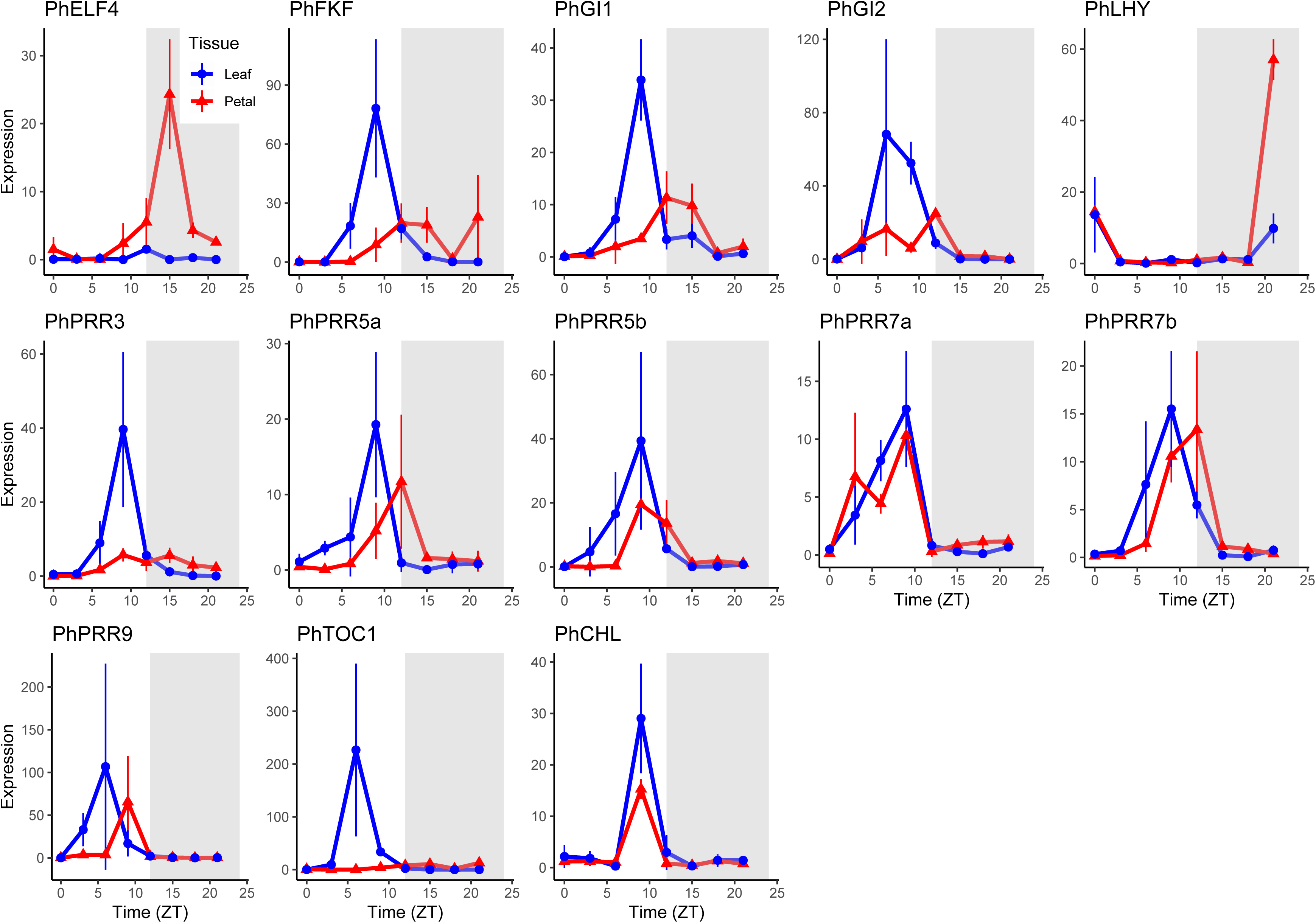
Daily changes in gene expression in petunia leaves and petals (12LD). Expression of clock genes in leaves (blue) and petals (red) under light:dark (LD 12 h : 12 h). Gene expression was analyzed by qPCR and normalized to *PhACT*. ZT0 (Zeitgeber time) denoting light on, and ZT12, light off; grey area indicates dark period. Results represent mean ± SD (n = 3).

We also found differences in amplitude between tissues. In general, amplitude of the clock genes was higher in leaves than in petals including *PhGI1, PhGI2, PhFKF* and the *PRR* genes *PhPRR9, PhPRR7b* and *PhTOC1*. The only gene showing larger amplitude in petals was *PhELF4* (Fig. 4, Supplementary Table S4). From all our observations we can conclude that the clock transcriptional structure differs in several ways between leaves and petals. First a robust rhythmic pattern was observed for all genes tested except *PhCHL* that was arrhythmic, *PhELF4* in leaves and *PhPRR7a* in petals. Most genes showed day phase in leaves and night phase in petals. Finally, the petal clock was somewhat dampened compared to leaves.

**Fig 4.**
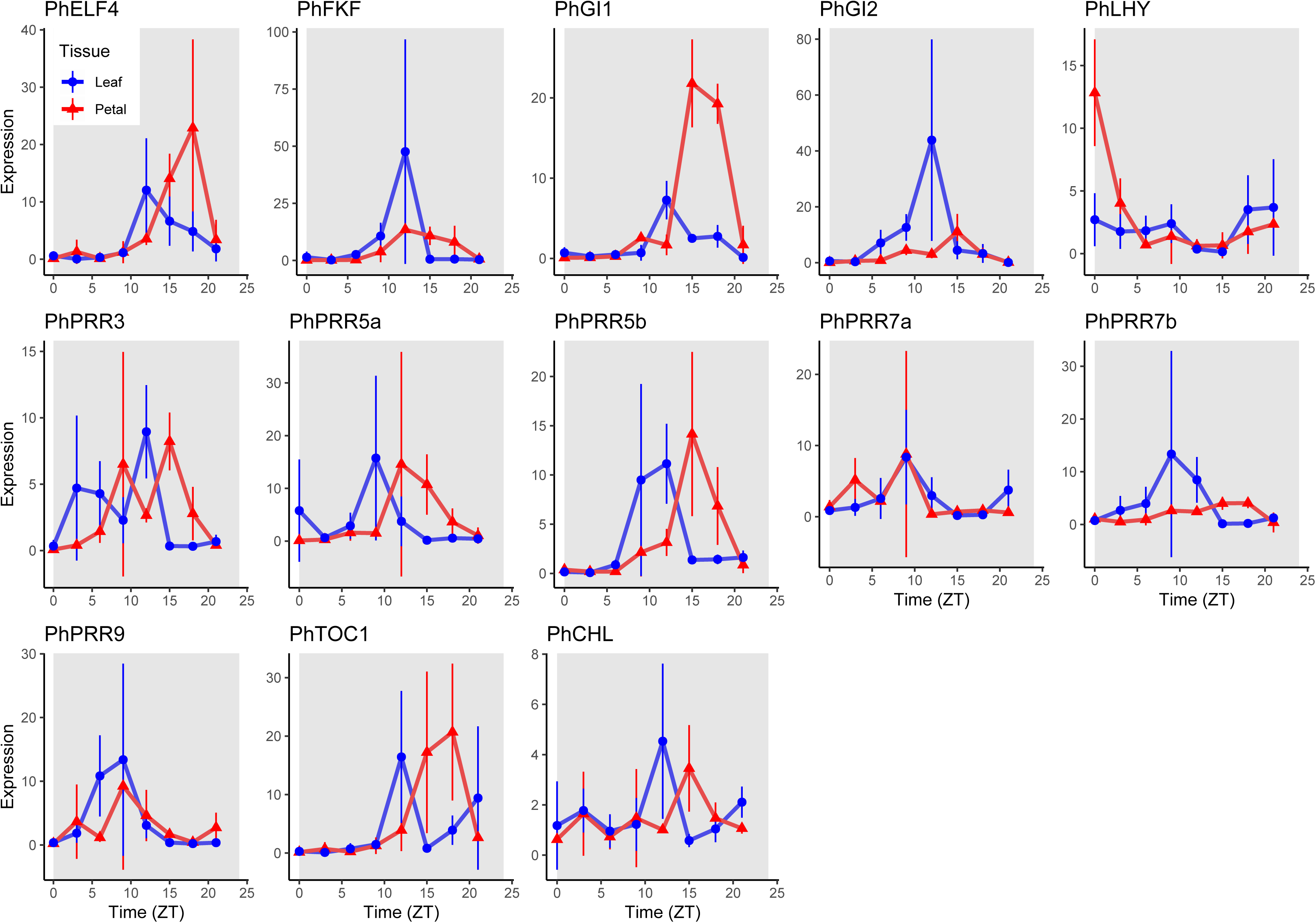
Daily changes in gene expression in petunia leaves and petals (12DD). Expression of clock genes in leaves (blue) and petals (red) under continuous dark. Gene expression was analyzed by qPCR and normalized to *PhACT*. Grey area indicates dark period, which includes subjective day (from ZT0, or Zeitgeber Time 0, to ZT12) and subjective night (from ZT12 to ZT24). Results represent mean ± SD (n = 3).

### The clock shows higher oscillation in petals than leaves under continuous dark

In order to study the entrainment of the petunia circadian clock to the light:dark cycle, petunia plants were transferred from light:dark (12LD) conditions to continuous darkness (12DD). Under constant darkness the genes *PhLHY* and *PhPRR7a* lost their significant oscillations in leaves (Table1). Interestingly, the gene *PhELF4* that was not rhythmic under LD conditions (Table 1) but displayed a robust oscillation in leaves under 12DD conditions. Finally, *PhPRR9* was not rhythmically expressed under a 12DD cycle in petals (Table 1). The rest of the genes analyzed maintained a rhythmic expression except for *PhCHL* that lacked a rhythm in any of the tissues or conditions analyzed, and *PhPRR7a* that was not rhythmic in petals.

We compared the expression between 12LD and 12DD in leaves (Fig. 4). We classified the clock genes in three groups either showing a delay in maximum expression between 1.5 and 7.5 hours (*PhPRR9, PhPRR5a, PhPRR5b, PhTOC1, PhGI1, PhGI2, PhFKF* and *PhCHL)* an advance: *PhPRR7b, PhPRR3* (1.5 hours) and *PhLHY* (18 hours) or a maintained maximum expression regardless of photoperiod (Table 1) (*PhPRR7a* and *PhELF4)*.

In petals, *PhPRR9, PhPRR7b, PhPRR5b, PhGI1, PhGI2* and *PhCHL* delayed their maximum expression between 1.5 and 10.5 hours. *PhPRR7a* and *PhLHY*, peaked 1.5 and 19.5 hours earlier, respectively. The last group included those genes that did not show differences in phase under 12LD or 12DD conditions: *PhPRR5a, PhPRR3, PhTOC1, PhELF4* and *PhFKF* (Table 1).

Altogether, *PhLHY* showed advanced expression under DD conditions while *PhGI1, PhGI2, PhPRR5b, PhPRR9* and *PhCHL* were delayed in both leaves and petals. The only gene that remained robust was *PhELF4*. Thus, these genes were homogenously affected by photoperiod. In contrast, *PhFKF, PhPRR5a, PhPRR7a, PhPRR7b, PhPRR3* and *PhTOC1* showed an organ specific change in phase in response to free running conditions (Table 1).

We compared the amplitude of clock genes in petunia leaves and petals under 12LD and 12DD. In leaves, we found that all genes showed a lower amplitude in continuous darkness except *PhELF4* displaying higher amplitude under 12DD (Supplementary Table S4). In petals the rhythmic expression dampened in *PhPRR9, PhPRR7a, PhPRR7b, PhPRR3, PhTOC1, PhGI1, PhGI2, PhFKF, PhCHL* and *PhLHY*. In contrast, the rhythm of *PhPRR5a, PhPRR5b* and *PhELF4* had higher amplitudes (Supplementary Table S4).

### Rhythmicity and photoperiod-sensitivity are tissue specific

An important paradigm in the analysis of circadian clock gene expression is the effect of free running conditions on the genes thought to have a circadian control (Somers *et al.*, 1998). We analyzed several parameters of circadian clock genes including phase, noise or amplitude in two tissues and light conditions using Harmonic ANOVA (Thaben and Westermark, 2016). These parameters resulted in a specific gene expression pattern that was compared in both tissues under LD and DD cycles (Table 2). We found that *PhELF4, PhLHY, PhPRR5a, PhPRR7a* and *PhPRR9* were stable regardless of the tissue or photoperiod (*p* > 0.05). In contrast, *PhFKF, PhGI1, PhPRR3* and *PhTOC1* showed a different expression pattern between leaves and petals under a 12LD cycle (*p* < 0.05). In contrast to LD conditions, under 12DD *PhGI1, PhPRR5b* and *PhPRR7b* were differentially expressed in leaf versus petal. When we compared leaves at 12LD versus 12DD, *PhGI1, PhGI2* and *PhTOC1* showed significant changes whereas in petals this group included *PhGI1, PhGI2, PhPRR5b, PhPRR7b and PhCHL* (Table 2).

**Table 2.**
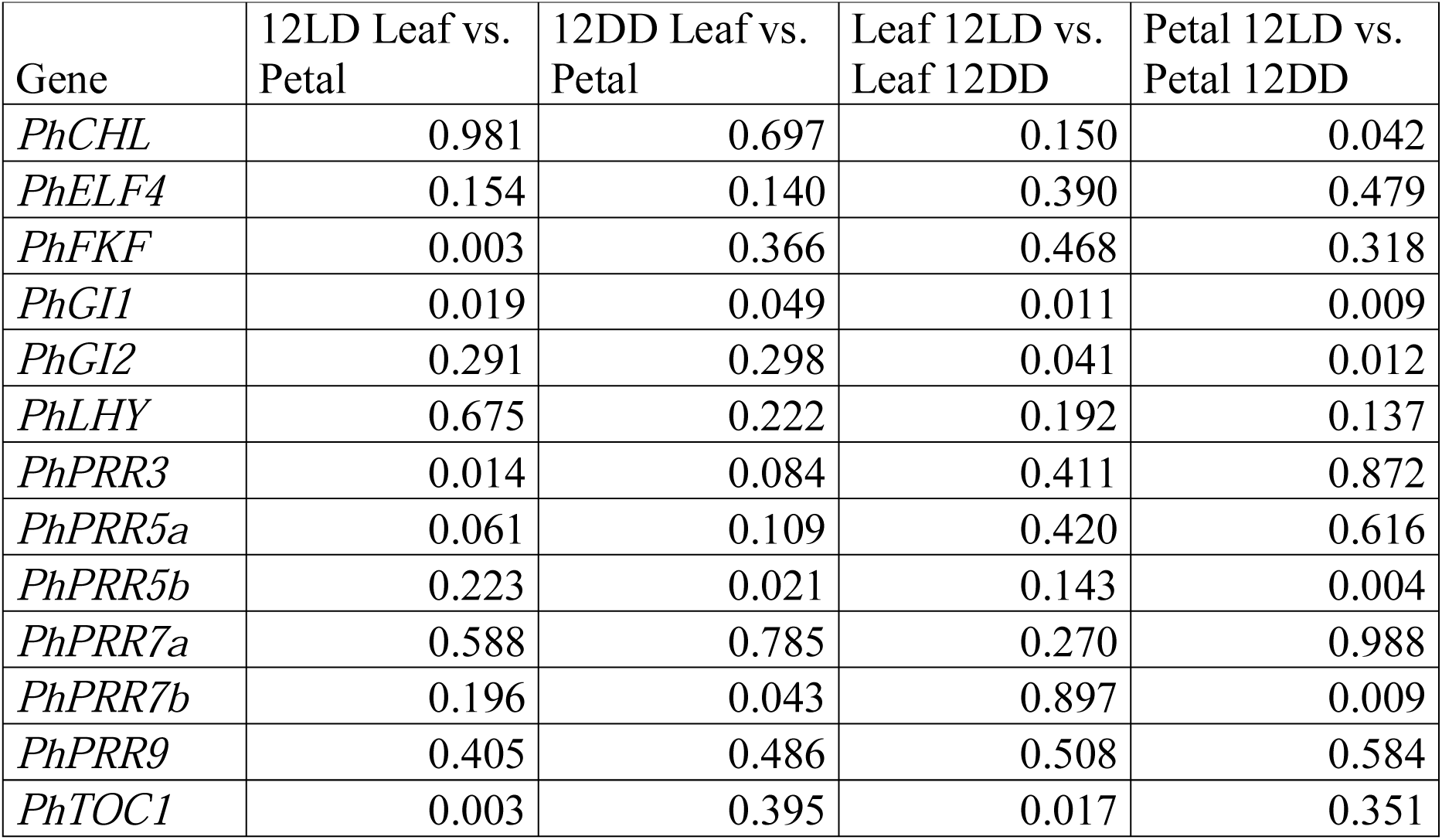
Analysis of differential gene expression in petunia leaves and petals under two light conditions: light:dark (12LD) and constant darkness (12DD). This analysis uses Harmonic ANOVA (HANOVA) to test differences. A *p* value < 0.05 indicated that the expression was significantly different between tissues (first and second column) or between light conditions (third and fourth column).

These results indicate that there are two sets of genes with different rhythms in leaves and petals and a group of stable genes comprising *PhELF4, PhLHY, PhPRR5a, PhPRR7a* and *PhPRR9*. Furthermore, the effect of photoperiod appeared to be organ-specific for those genes that showed significant changes.

### Transcriptional noise is gene and tissue specific

Although gene expression quantities were determined for the same set of mRNA extractions, the degree of significance in terms of gene expression levels was not always as expected based on average expressions. This indicated that some genes had robust expression levels while others appeared to be very variable. In order to quantify the dispersion of data, we plotted the normalized Ct values for all genes, dividing the Ct of the clock gene by the Ct of the reference gene *PhACT* (Fig. 5, Fig. 6) and calculated the coefficient of variation (CV) for all time points (Supplementary Table S5). We found that the data dispersion was very different between genes, tissues and light conditions. The gene with the maximum transcriptional noise was *PhLHY* in petals at ZT0 and 12LD (CV 24.81) while *PhPRR7a* in leaves showed the lowest at ZT0 and 12LD (CV 0.56) (Supplementary Table S5). In addition, transcriptional noise seemed to change during the day. In leaves under a light:dark cycle, the highest noise was found at ZT9 (average CV 9.19) and the lowest, at ZT18 (average CV 4.35). In contrast, in petals, the maximum noise was at ZT0 (average CV 10.33) and the minimum, at ZT12 (average CV 3.44) (Fig. 5, Supplemental Table S5). Under constant darkness, this pattern varied. Leaves, displayed the highest CV at ZT12 (average CV 7.89) and the lowest, at ZT0 (average CV 4.34). Petals showed the maximum transcriptional noise at ZT9 (average CV 9.41) and the minimum at ZT12 (average CV 3.31) (Fig. 6, Supplementary Table S5).

**Fig 5.**
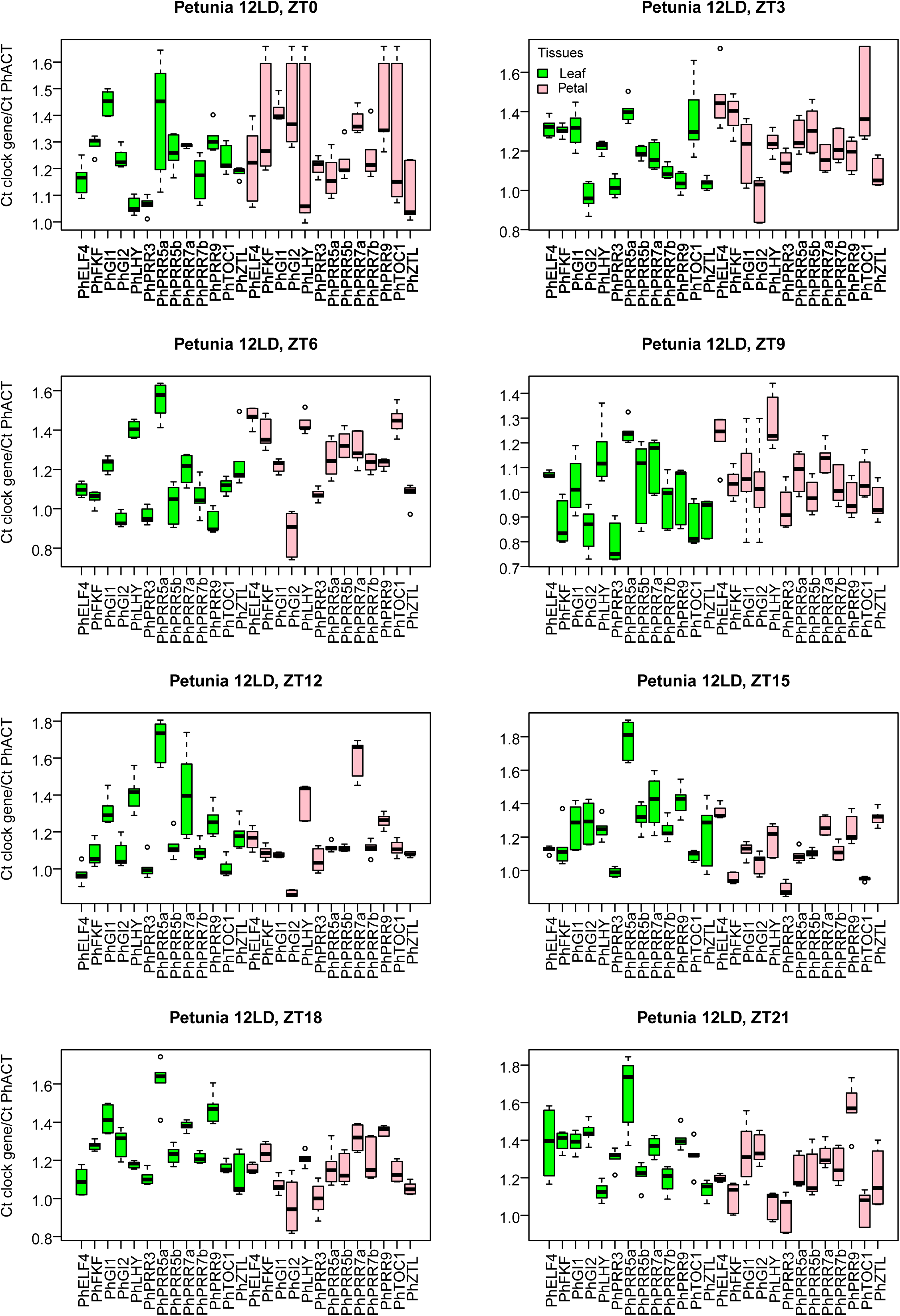
Boxplot of cycle threshold values (Ct) for petunia clock genes normalized with *PhACT* (Ct of clock gene divided by Ct of *PhACT*) in leaves (green) and petals (pink) under constant darkness (12LD) at eight time points, from ZT0 to ZT21.

**Fig 6.**
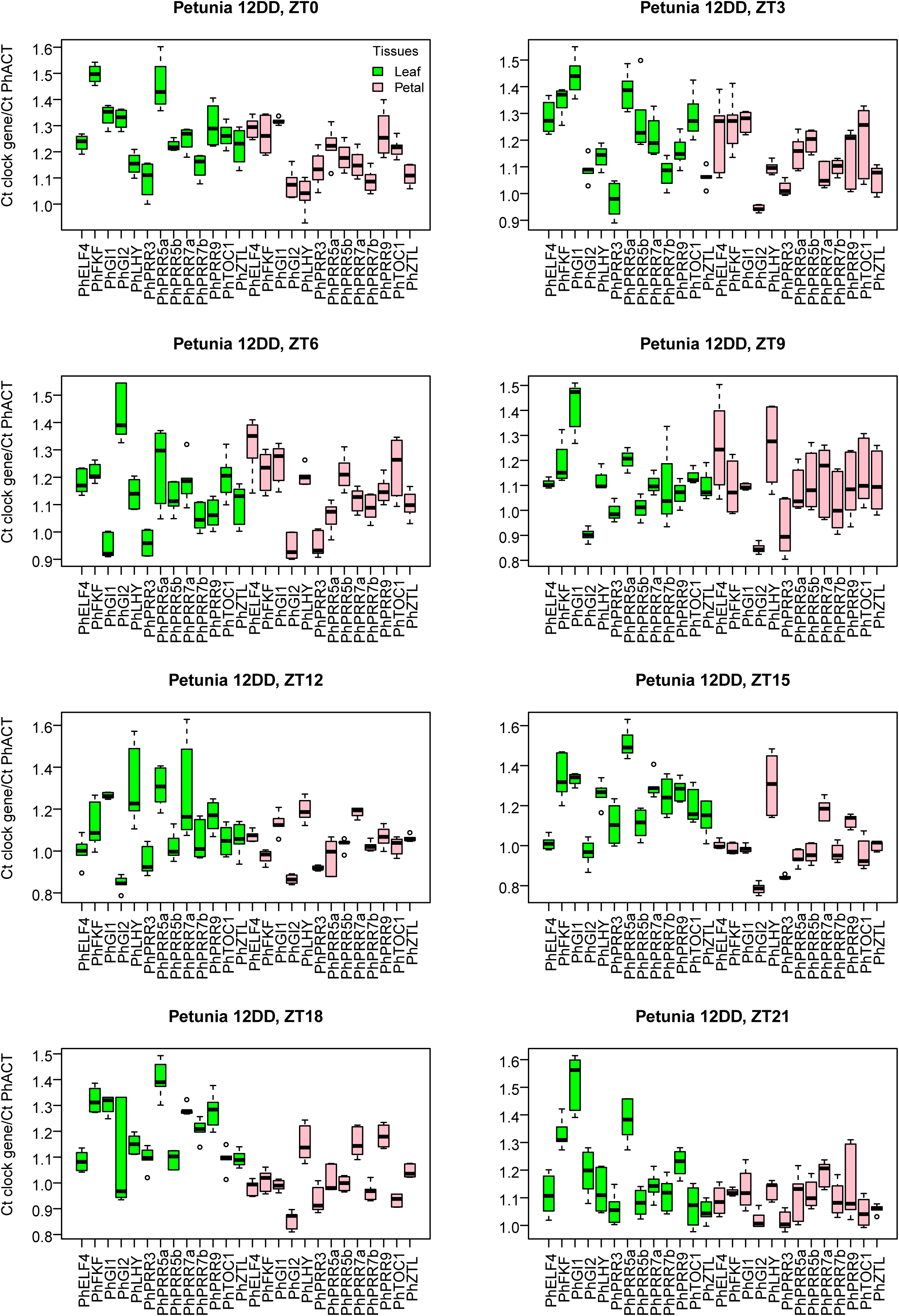
Boxplot of cycle threshold values (Ct) for petunia clock genes normalized with *PhACT* (Ct of clock gene divided by Ct of *PhACT*) in leaves (green) and petals (pink) under constant darkness (12DD), at eight time points, from ZT0 to ZT21.

We can conclude that subjective time ZT0 i.e. when lights are turned on, displayed the lowest transcriptional noise in leaves and the highest in petals. When day advanced, noise increased in leaves that showed its maximum at ZT9 with opposite behavior in petals that had its lowest level of noise at ZT12 i.e. when lights were turned off. Under free running conditions, the same pattern was found as the lowest and highest noise for leaves coincided with early and late day respectively, while in petals transcriptional noise was low in the subjective night and higher noise was found at subjective time ZT9. This indicates that an endogenous component governs transcriptional noise of the clock genes, which also differs in leaves and petals.

## Discussion

### The petunia clock gene show structural evolutionary changes

The evolution of the plant circadian clock is considered an important driver of adaptation in a variety of plants including tomato, *Opuntia ficus-indica* or barley (Mallona *et al.*, 2011*a*; Zakhrabekova *et al.*, 2012; Müller *et al.*, 2016; Müller *et al.*, 2018). The plant clock is an important coordinator of primary and secondary metabolism in plants. It defines the timing of floral scent emission in a variety of plants including *Petunia* or *Nicotiana attenuatta* (Fenske *et al.*, 2015; Yon *et al.*, 2015; Terry *et al.*, 2019). The plant circadian clock appears to have a specific transcriptional structure in different tissues such as leaves, pods, seeds, or roots (Thain *et al.*, 2002; James *et al.*, 2008; Bordage *et al.*, 2016; Weiss *et al.*, 2018). As the transcriptional structure of the clock in petal is currently unknown, we used *Petunia hybrida* to perform a detailed analysis. We have characterized the structural changes in *PhPRR5a, PhPRR5b, PhPRR7a, PhPRR7b, PhGI1* and *PhGI2* and the transcriptional structure of the petunia circadian clock in petals and leaves, using standard growth and free running conditions of continuous darkness.

The complete genome paleohexaploidization of petunia, found in the Solanaceae group (Bombarely *et al.*, 2016) is reflected in the retaining of several clock genes as duplications that are found as single copy genes in Arabidopsis and other species. These include *PhPRR5a, PhPRR5b, PhPRR7a, PhPRR7b, PhGI1* and *PhGI2.* Other genes that are found as single copy include *PhLHY, PhPRR9, PhPRR3, PhTOC1, PhFKF* and *PhCHL*. Interestingly genes found as single copy in petunia such as *PhTOC1, PhPRR9* and *PhPRR3* are found as single copy in most Solanaceae except for *N. benthamiana* that appears to have two copies of each gene (Fig 1). Two of the petunia paralogs *PhPRR7a, PinfS6PRR7a* and *PhPRR5a* and *PinfS6PRR5a* cluster between Arabidopsis and the rest of the Solanaceae genes. In contrast the single copy genes *TOC1, PRR3* and *PRR9* are found as a subclade for all the Solanaceae together including *Petunia*. This indicates that there has been a loss of *PRR5* and *PRR7* paralogs in the Solanaceae that have a single copy gene, while *Petunia* has retained the older copy closer to the Arabidopsis, *Vitis vinifera* and *Amborella trichopoda* genes. The additional changes observed in the number of exons indicate a specific evolution of one paralog. Indeed, *AtPRR5* has six exons whereas *AtPRR7* presents nine exons (AT5G24470.1 and AT5G02810, consulted in TAIR database) while *PhPRR5a* and *PhPRR7b* present 7 exons whereas *PhPRR5b* and *PhPRR7a* have 8 exons, indicating possible sub or neofunctionalization of these paralogs (see below).

We found two domains, REG and CCT in all analyzed TOC1, PRR3, PRR5 and PRR9 sequences. In contrast, the CCT domain was absent in most PRR7 paralogs in *Capsicum spp*., *Petunia spp*., *Solanum spp.* and *Nicotiana spp.* Interestingly, we only found the CCT domain in PhPRR7a, which shared more similarities in the amino acids sequence with AtPRR7. The lack of CCT domains in Solanaceae but not in the related Convolvulaceae family suggests that this event occurred in the early history of Solanaceae. In addition, this alteration, which has been has been described in PRR orthologs in crops such as rice and soybean, can modify growth and flowering time (Lenser and Theißen, 2013; Li *et al.*, 2019*a*). This may result in a specific clock in the Solanaceae family.

The gene *GI* appeared in flowering plants and is absent in mosses or picoalgae (Linde *et al.*, 2017). In the Solanaceae we found two to three copies, and in *Petunia hybrida*, there are significant differences in the coding region between *PhGI1* and *PhGI2* suggesting a diversification of functions. Furthermore, the amino acid differences between *P. axillaris* and *P. inflata* indicate species specific changes in this master regulator that maybe related to the differing environmental niches where both species grow.

We used the predicted protein sequences to infer the domain structure of GIGANTEA. Although a previous study describes that *GI* encodes a protein with six transmembrane domains (Park *et al.*, 1999), the biochemical functions of GI are not understood. Yeast two hybrid experiments performed with the Arabidopsis GI protein show that the N-terminal domain interacts with FKF1 (Sawa *et al.*, 2007), while the complete protein shows interactions with the CYCLING DOF FACTOR6 protein (Krahmer *et al.*, 2019). As the differences in protein structure found between PhGI1 and PhGI2 do not match well known domains we cannot understand their functional differences. Nevertheless, the PinfS6GI3 does lack the N terminus required for interactions with FKF1 and ZTL in Arabidopsis.

### Daily expression of petunia clock genes is tissue specific

The current transcriptional model of the plant circadian clock is largely based on the expression of genes in the Arabidopsis hypocotyls and leaves (Staiger *et al.*, 2013). It includes the morning, midday or core and the evening loops. During the morning, the genes *CCA1* and *LHY* repress the evening genes *GI* and *TOC1* and activate *PRR9* and *PRR7*. At the same time, *TOC1* acts repressing *GI* and *PRR9* but activating *CCA1/LHY*. On the other hand, GI stabilizes ZTL that is a *TOC1* repressor (Pokhilko *et al.*, 2010).

Previous studies have revealed that the circadian clock is tissue-specific (Thain *et al.*, 2002; Endo *et al.*, 2014; Bordage *et al.*, 2016). Differential expression of clock genes has been reported in several tissues including seeds, roots, leaves, stems and flowers at several developmental stages in different plant species such as bamboo (Dutta *et al.*, 2018), radish (Wang *et al.*, 2017) or daisy (Fu *et al.*, 2014). The present study has covered several clock genes, including *GI* and *PRRs* paralogs, in petunia leaves and petals and our results are consistent with the existence of organ-specific biological clocks in plants.

### The expression of clock genes differs between paralogs

Changes in gene expression concerning timing, quantity and rhythm may hint at possible subfunctionalization or neofunctionalization of duplicated clock genes. We found that *PhGI1, PhGI2, PhPRR7b* and *PhPRR5b* had similar expression patterns to those previously described in other plants in leaves (Fowler *et al.*, 1999; Matsushika *et al.*, 2000; Marcolino-Gomes *et al.*, 2014). In contrast, *PhPRR5a* and *PhPRR7a* that were the closest paralogs to the rest of the species, showed modified expression patterns. *PhPRR5a* and *PhPRR7a* showed an advanced phase, peaking before their respective paralogs, *PhPRR5b* and *PhPRR7b*. Interestingly, in petals, *PhPRR7a* displayed a profile similar to the canonical *AtPRR7*. Moreover, the paralogs *PhGI1, PhGI2, PhPRR5a, PhPRR5b* and *PhPRR7b* delayed their maxima to the dark period.

### Leaves and petals have different clock coordination

In the present work we identified significant oscillations in gene expression using the JTK_CYCLE algorithm, a non-parametric method which also provided measures of phase and period (Hughes *et al.*, 2010). As mentioned above, most analyzed genes displayed a robust rhythm. Second, we performed an HANOVA test and we found genes that displayed a differential expression pattern, comparing tissues and light conditions. The core clock genes *LHY* and *TOC1* are found in basal picoeukaryotes, mosses, *Marchantia polymorpha* and all higher plants (Corellou *et al.*, 2009; Holm *et al.*, 2010; Linde *et al.*, 2017). We found that *PhLHY* and *PhPRR9* did not show any statistical differences regardless the tissue or light cycle. In contrast, *PhTOC1* expression pattern differed between leaves and petals. This indicates a basal change in the clock coordination between both tissues. This scenario maybe further supported by the significant changes found for *PhFKF, PhPRR3*, and *PhGI1* between tissues. Finally, *PhGI1*, a gene found only in flowering plants showed significant changes between tissues and photoperiods indicating that it may play a role in the coordination between development and environmental signals.

### Photoperiod sensitivity is organ-specific

The effect of day length on biological clocks has been widely studied. For example, floral transition is controlled by *CONSTANS (CO)* and *FLOWERING LOCUS T (FT)* genes which are regulated by the circadian clock, including *ELF3, ELF4, GI, LHY, PRRs* and *ZTL* genes (Samach *et al.*, 2000; Suárez-López *et al.*, 2001; Valverde *et al.*, 2004). These genes are capable to integrate environmental cues, mainly day length, but also temperature. Clock genes are therefore sensitive to ambient changes resulting in an adaptive advantage (Dodd *et al.*, 2005). The present study revealed that a constant dark regime induced phase-shift even in the first 24h. Most analyzed genes tended to delay their maximum expression, especially in leaves. Only *PhLHY* advanced its phase both in leaves and petals. Interestingly *PhLHY* lost its rhythmic expression in leaves but it persisted in petals, similar to previous studies (Fenske *et al.*, 2015). Other genes, *PhPRR7a* (in leaves) and *PhPRR9* (in petals), did not retain their rhythmicity, suggesting that the integration of environmental cues and phototransduction varies depending on the tissue. This is consistent with previous studies, that have reported the effect of light on organ-specific circadian clocks and photoperiodic sensitivity (Shimizu *et al.*, 2015; Bordage *et al.*, 2016).

Constant dark also had an effect on oscillations, which in general tended to decrease in most analyzed genes in leaves and petals. Similar results have been reported in other plants species: *LHY/CCA1, ELF4, GI* and *TOC1* gene expression dampens under constant light or constant dark conditions in Arabidopsis (Wang and Tobin, 1998; Park *et al.*, 1999; Liew *et al.*, 2014; Fenske *et al.*, 2015). Loss of circadian rhythmicity could be key and be involved in responses to environmental changes, such as seasonal dormancy during winter in Japanese cedar or chestnut (Ramos *et al.*, 2005; Nose and Watanabe, 2014).

### Transcriptional noise is tissue-specific and depends on the photoperiod

One of the main features of the transcriptional structure of circadian clocks is the capacity to integrate noisy environmental signals and internal transcriptional variation (Hogenesch and Ueda, 2011). The robustness of circadian oscillation is related to the number of mRNA molecules, interactions and complex formation, and it is stabilized by the entrainment to the light:dark cycle (Gonze *et al.*, 2002).

In the present work we found that molecular noise differed in leaves and petals and it was influenced by the time of the day. While in leaves highest stability appeared at the beginning of the subjective day, petals displayed the lowest stability. This was also noticeable when plants were transferred to continuous darkness. Interestingly, the time point with the highest transcriptional noise shifted both in leaves and petals. The lowest stability advanced in petals, and delayed in leaves. Furthermore, the increased transcriptional robustness early in the day in leaves, and in the late day-early night in petals, coincide with the major functional changes in both tissues, initiation of photosynthesis and scent emission. As noise increases thereafter in both tissues, it could be that funneling transcriptional noise into robustness at certain times of the day may have biological implications to achieve consistent outputs. However, the molecular function, if any, is not understood as this is the first report of this phenomenon.

Taken together the differential transcriptional structure and response to light, we conclude that the circadian clock in leaves and petals show substantial differences, that may reflect the underlying function in controlling photosynthesis and secondary metabolism in both tissues. The functional differences between leaves and petals may rely in part on a circadian clock reprogramming during flower development.

## Supporting information

**Fig. S1**. Melt or dissociation curve analysis of petunia genes.

**Fig. S2**. Exon-intron structure of *Petunia axillaris* (PaxiN) *PRR5* and *PRR7* genes.

**Fig. S3**. (A) Domain structure of PRRs proteins.

**Fig. S4**. Local alignment of GIGANTEA proteins.

**Table S1**. PSEUDO-RESPONSE REGULATORs (PRRs) protein accessions used in the phylogenetic reconstruction and for the annotation of protein sequences.

**Table S2**. GIGANTEA (GI) protein accessions used in the phylogenetic reconstruction.

**Table S3.** Primers used for qPCR.

**Table S4**. Rhythmic analysis of transcriptional data.

**Table S5**: Coefficient of variation, gene expressions.

## Authors’ contributions

MIT, MCS, and MEC performed the experimental work; MIT, JW and MEC designed the research programme; JW and MEC secured funds; MIT, JW and MEC wrote the first draft of the manuscript and all authors commented and corrected the final manuscript.

## Acknowledgements

This work was developed under projects Fundación Séneca 19398/PI/14, MICINN-FEDER BFU-2013-45148-R and BFU-2017-88300-C2-1-R.

## Abbreviations

CCT: CONSTANS, CONSTANS-like, and TIMING OF CAB EXPRESSION 1 domain
Ct: Cycle threshold
PaxiN: Petunia axillaris
PhACT: ACTIN
PhELF4: EARLY FLOWERING 4
PhFKF: FLAVIN-BINDING KELCH REPEAT F-BOX
PhGI1: GIGANTEA 1
PhGI2: GIGANTEA 2
PhLHY: LATE ELONGATED HYPOCOTYL
PhPRRs (PhPRR3, PhPRR5a, PhPRR5b, PhPRR7a, PhPRR7b and PhPRR9): PSEUDO-RESPONSE REGULATORS
PhTOC1: TIMING OF CAB EXPRESSION 1
PhCHL: CHANEL (ZEITLUPE)
Ph: Petunia hybrida
PinfS6: Petunia inflata
REG: Response regulatory domain
ZT: Zeitgeber time.

